# Environmental and anthropogenic drivers of *Culex pipiens* biotype *pipiens* overwintering phenology

**DOI:** 10.64898/2026.11.23.768647

**Authors:** Carmen Villacañas de Castro, Ellen Kiel, Renke Lühken, Felix Gregor Sauer

**Affiliations:** Bernhard Nocht Institute for Tropical Medicine, Hamburg, Germany; Carl von Ossietzky University, Oldenburg, Germany

**Keywords:** *Culex pipiens* biotype *pipiens*, phenology, overwintering, land use, artificial light at night, urban heat island

## Abstract

**Background:** *Culex pipiens* biotype *pipiens* (*Cx. p. pipiens*) is the primary vector of West Nile and Usutu viruses in Europe and overwinters as adult females in shelters, such as cellars. However, the localised environmental and anthropogenic drivers regulating adult behavioural plasticity and the timing of entry into and exit from these overwintering sites remain poorly understood.

**Methods:** Over two consecutive winters (2023–2025) in Northwestern Germany, we monitored the overwintering period of *Cx. p. pipiens* across an urban-rural gradient using custom interception traps at the entry points (e.g. windows) of overwintering sites (e.g. cellars). We modelled the days of the year at which 10%, 50%, and 90% of cumulative populations entered or exited shelters (Q_10_, Q_50_, Q_90_), against satellite-derived land use and artificial light at night (ALAN), environmental temperature, indoor temperature and relative humidity, cumulative degree-days (CDD), and wing size.

**Results:** In autumn, early entry (Q_10_) was delayed by 24.1±7.7 days in semi-urban allotment gardens compared to rural sites. At late entry (Q_90_), land use effects disappeared, and non-linear deviations from seasonal mean environmental temperatures shaped entry across land uses, while increasing ALAN delayed entry (0.64 days per unit increase). Entry probability increased with cumulative cooling degree-days below 18 °C. In spring, exit timing was independent of land use but driven by the indoor microclimate. Early exit (Q_10_) was advanced by higher minimum indoor temperatures (2.35 days per 1 °C) and higher minimum relative humidity (0.57 days per 1%), with additive land use effects driving earlier emergence in urban than in rural sites. Conversely, higher minimum relative humidity delayed late exit (Q_90_). Spring exit was best predicted by CDD above 0 °C. Finally, larger wing centroid size did not affect overwintering survival but delayed spring exit in *Cx. p. pipiens*.

**Conclusion:** Adult *Cx. p. pipiens* exhibit behavioural plasticity, shortening their overwintering period in human-altered landscapes. Delayed autumn entry, paired with earlier spring exit driven by urban warming and low thermal thresholds, extends seasonal vector activity. Integrating these complex anthropogenic and microclimatic drivers into surveillance frameworks is vital for predicting early-season pathogen persistence and optimising targeted interventions.

## Background

The *Culex pipiens* complex is a group of widely distributed mosquito species, known as vectors for arboviruses of medical and veterinary relevance, including West Nile virus, Usutu virus, or Sindbis virus [1, 2, 3, 4]. In Europe, *Cx. pipiens sensu stricto* (s.s.) can be differentiated into two morphologically indistinguishable biotypes: biotype *pipiens*, which is anautogenous and overwinters in a state of diapause; and biotype *molestus*, which is stenogamous, subterranean, and does not undergo overwintering diapause [1]. Among these, *Cx. pipiens* biotype *pipiens* (*Cx. p. pipiens*) is the predominant member and exhibits a broad host-feeding pattern [5], facilitating virus circulation among avian hosts and acting as a bridge vector to humans and other mammals [6, 7, 8]. Understanding the ecology and seasonal dynamics of this species is essential for predicting arbovirus transmission risk.

In temperate climates, *Cx. p. pipiens* overwinters as inseminated adult females, whereas the males die during autumn. The overwintering ability of the females is therefore a central component of mosquito population dynamics and ensures local maintenance of mosquito populations [9, 10]. In addition, the detection of different pathogens in overwintering *Culex* mosquitoes demonstrated that they can serve as reservoirs for arboviruses including Sindbis [11], Usutu [12, 13] and West Nile virus [14, 15, 16], thus contributing to the arbovirus persistence in temperate regions. As reviewed by DL Denlinger and PA Armbruster [9], females entering diapause display a distinctive phenotype, characterised by reproductive arrest, increased sugar feeding, and enhanced lipid accumulation. Their facultative diapause is induced by environmental cues, particularly shortening photoperiods and decreasing temperatures experienced during the immature stages at the end of summer [9, 17].

Field studies in Central Europe have shown that *Cx. p. pipiens* females overwinter in anthropogenic subterranean sites, such as cellars, tunnels, or other protected constructions [11, 15, 18], as well as in natural shelters like caves or animal burrows [13, 19]. These overwintering sites are usually characterised by a stable microclimate with low light exposure, cool temperatures, and moderate to high humidity [18, 20, 21]. The survival of overwintering *Cx. p. pipiens* is strongly affected by microclimate. Stable, moderately humid sites support higher overwintering densities and increased survival rates [18]. J Rozsypal, M Moos, I Rudolf and V Košťál [22] found that the suitable overwintering sites for *Cx. p. pipiens* had limited air movements, and temperatures between 2 °C and 6 °C, whereas sub-zero temperatures increased mortality after only a few days. These studies suggest a non-random habitat selection, with mosquitoes preferentially aggregating in subterranean structures that buffer environmental variability. Even small differences in microclimate may alter the rate of lipid reserve depletion, eventually impacting mosquito survival, post-winter development thresholds, or phenology [9, 23, 24].

However, the relative influence of environmental and anthropogenic drivers that shape the onset and termination of diapause is still poorly understood. Urbanisation may alter overwintering dynamics by modifying site availability and local microclimatic conditions [25, 26]. In addition, artificial light at night (ALAN) has been shown to interfere with circadian and seasonal activity patterns across multiple insect species [27, 28, 29, 30], and alters diapause timing in several mosquito vectors [31, 32, 33, 34].

In addition to phenological shifts caused by anthropogenic changes of the cues inducing diapause, adult mosquitoes may display behavioural plasticity. While the diapause phenotype is induced by photoperiodic and thermal cues during the immature larval stages [9], the specific timing of entry to and exit from the overwintering shelters may be further modulated by adult behaviour. Diapausing females may dynamically adjust their activity, sugar feeding, and shelter seeking behaviours in response to local environmental cues [9]. Consequently, a significant behavioural lag can occur between diapause induction/deduction and entry/exit of the overwintering site, as environmental or anthropogenic drivers alter local conditions [25, 33].

To address these knowledge gaps, this study investigates the overwintering behaviour of *Cx. p. pipiens* in Germany, focusing on environmental and anthropogenic drivers across an urban gradient. Specifically, we aim to (i) quantify overwintering phenology, including both entry into and exit from overwintering sites, (ii) examine the influence of ALAN on mosquito activity patterns, and (iii) evaluate how climatic variables (e.g., temperature and relative humidity) modulate adult behaviour and affect entry and exit timing. By integrating field observations across contrasting land use types and two consecutive winters, this study aims to clarify how environmental conditions regulate overwintering behaviour, improving our understanding of the drivers shaping overwintering dynamics in a crucial vector species.

## Materials and Methods

### Selection of the study sites

This study was conducted in Northern Germany in cities of Oldenburg (Lower Saxony) and Bremen between autumn 2023 and spring 2025. Bremen, one of Germany’s larger cities (600,000 inhabitants), represents a high-density urban environment, whereas Oldenburg, a smaller city (180,000 inhabitants), represents a lower-density, mid-sized urban environment. Both cities exhibit high habitat heterogeneity and diverse land use patterns, making them well-suited for investigating the effects of varying microclimatic conditions and the influence of ALAN on the overwintering dynamics of mosquitoes.

Potential overwintering sites located in cellars, basements, or sheds were identified through direct engagement with private property owners, local residents, and personal networks. Once a site was confirmed to be suitable for our study (unheated structures with external access/windows, and/or confirmed presence during winter) its geographic coordinates were recorded using a GPS (GARMIN etrex 20x, Garmin Deutschland, Garching, Germany) extracted at the exact point location. These coordinates were then used to extract macrohabitat variables. Land use at each sampling location was classified into three broad categories: urban, semi-urban, and rural. Classification was based on the CORINE Land Cover 2018 (CLC2018) raster dataset. The data were processed using the open-source geographic information system software QGIS QGIS Development Team, 2024. CLC land use classes were aggregated as follows:

• urban: continuous and discontinuous urban fabric, industrial or commercial units

• semi-urban: sport and leisure facilities, predominantly represented by *Kleingartenvereine* (allotment garden complexes commonly found in German cities)

• rural: pastures and non-irrigated arable land

ALAN was quantified using satellite-derived radiance data from the Light Pollution Map (https://www.lightpollutionmap.info), which is based on nighttime light observations from the Visible Infrared Imaging Radiometer Suite (VIIRS). Radiance values (nW·cm⁻²·sr⁻¹) were extracted based on the GPS coordinates of each sampling site location and used as a proxy for local light pollution intensity.

Our goal was to replicate the study across two cities over two consecutive winters, including three land use categories with five overwintering sites per category. The distribution of sampling sites among land use categories varied slightly between study winters (Table 1) due to changes in property access and site availability. In winter 2023–2024, sampling sites in Bremen comprised 6 urban, 6 semi-urban, and 4 rural sites, while Oldenburg included 4 urban, 2 semi-urban, and 4 rural sites. In winter 2024–2025, Bremen included 5 sites per category, whereas Oldenburg comprised 5 urban and 5 rural sites.

**Table 1.** Geographic coordinates and land use of 2023/2024 and 2024/2025 monitored overwintering sites in Bremen and Oldenburg.

| 2023/2024 |  |  |  |  | 2024/2025 |  |  |  |  |
|---|---|---|---|---|---|---|---|---|---|
| ID | City | Longitude | Latitude | Land use | ID | City | Longitude | Latitude | Land use |
| HB01_2023 | Bremen | 53.129013 | 8.628644 | rural | HB01_2024 | Bremen | 53.129013 | 8.628644 | rural |
| HB02_2023 | Bremen | 53.158633 | 8.831032 | rural | HB02_2024 | Bremen | 53.118551 | 8.816142 | rural |
| HB03_2023 | Bremen | 53.072630 | 8.909335 | urban | HB03_2024 | Bremen | 53.072630 | 8.909335 | urban |
| HB04_2023 | Bremen | 53.064801 | 8.807365 | urban | HB04_2024 | Bremen | 53.063214 | 8.792509 | urban |
| HB05_2023 | Bremen | 53.077520 | 8.767430 | semi-urban | HB05_2024 | Bremen | 53.077520 | 8.767430 | semi-urban |
| HB06_2023 | Bremen | 53.173997 | 8.892559 | rural | HB06_2024 | Bremen | 53.173997 | 8.892559 | rural |
| HB07_2023 | Bremen | 53.063875 | 8.816807 | semi-urban | HB07_2024 | Bremen | 53.118842 | 8.639705 | rural |
| HB08_2023 | Bremen | 53.062143 | 8.834048 | semi-urban | HB08_2024 | Bremen | 53.070281 | 8.756870 | semi-urban |
| HB09_2023 | Bremen | 53.101825 | 8.780166 | urban | HB09_2024 | Bremen | 53.101825 | 8.780166 | urban |
| HB10_2023 | Bremen | 53.093572 | 8.800534 | urban | HB10_2024 | Bremen | 53.093572 | 8.800534 | urban |
| HB11_2023 | Bremen | 53.091788 | 8.848532 | urban | HB11_2024 | Bremen | 53.054266 | 8.773119 | semi-urban |
| HB12_2023 | Bremen | 53.066005 | 8.791111 | urban | HB12_2024 | Bremen | 53.066005 | 8.791111 | urban |
| HB13_2023 | Bremen | 53.129152 | 8.867286 | rural | HB13_2024 | Bremen | 53.129152 | 8.867286 | rural |
| HB14_2023 | Bremen | 53.084374 | 8.755716 | semi-urban | HB14_2024 | Bremen | 53.084374 | 8.755716 | semi-urban |
| HB15_2023 | Bremen | 53.087931 | 8.758330 | semi-urban | HB15_2024 | Bremen | 53.087931 | 8.758330 | semi-urban |
| HB16_2023 | Bremen | 53.065231 | 8.825051 | semi-urban | - | - | - | - | - |
| OL01_2023 | Oldenburg | 53.317089 | 8.250678 | rural | OL01_2024 | Oldenburg | 53.317089 | 8.250678 | rural |
| OL02_2023 | Oldenburg | 53.229858 | 8.318852 | rural | OL02_2024 | Oldenburg | 53.229858 | 8.318852 | rural |
| OL03_2023 | Oldenburg | 53.216070 | 7.748690 | rural | OL03_2024 | Oldenburg | 53.120898 | 8.157297 | rural |
| OL04_2023 | Oldenburg | 53.223173 | 7.735661 | rural | OL04_2024 | Oldenburg | 53.167153 | 8.160623 | urban |
| OL05_2023 | Oldenburg | 53.141648 | 8.191352 | urban | OL05_2024 | Oldenburg | 53.141648 | 8.191352 | urban |
| OL06_2023 | Oldenburg | 53.146477 | 8.180513 | urban | OL06_2024 | Oldenburg | 53.145997 | 8.184139 | urban |
| OL07_2023 | Oldenburg | 53.155901 | 8.235612 | urban | OL07_2024 | Oldenburg | 53.155392 | 8.234816 | urban |
| OL08_2023 | Oldenburg | 53.139074 | 8.139082 | urban | OL08_2024 | Oldenburg | 53.139074 | 8.139082 | urban |
| OL09_2023 | Oldenburg | 53.154040 | 8.245695 | semi-urban | OL09_2024 | Oldenburg | 53.202801 | 8.339346 | rural |
| OL10_2023 | Oldenburg | 53.163139 | 8.183392 | semi-urban | OL10_2024 | Oldenburg | 53.145043 | 8.139074 | rural |

### Phase 1: Entry into overwintering sites

During autumn, we monitored mosquito entry into the overwintering sites using interception traps. These were installed at the main entry points to each site (specifically, windows across all sites, except for one ventilation shaft during the first winter) in early September 2023 and mid-August 2024. Each entry point was covered with a commercial mosquito net that was fixed to the surrounding walls to fully seal the opening. A circular opening was cut in the centre of the net and fitted with hook-and-loop fastener strips to attach a custom-made mosquito trap (Figure 1a). The trap consisted of a plastic bottle neck used as a funnel, connected to a cylinder constructed with chicken wire and mosquito netting, and secured using cable ties and hot glue. A catch bag (18 cm × 18 cm; Biogents, Regensburg, Germany) was attached to the distal end of the trap to retain captured mosquitoes.

**Figure 1.**
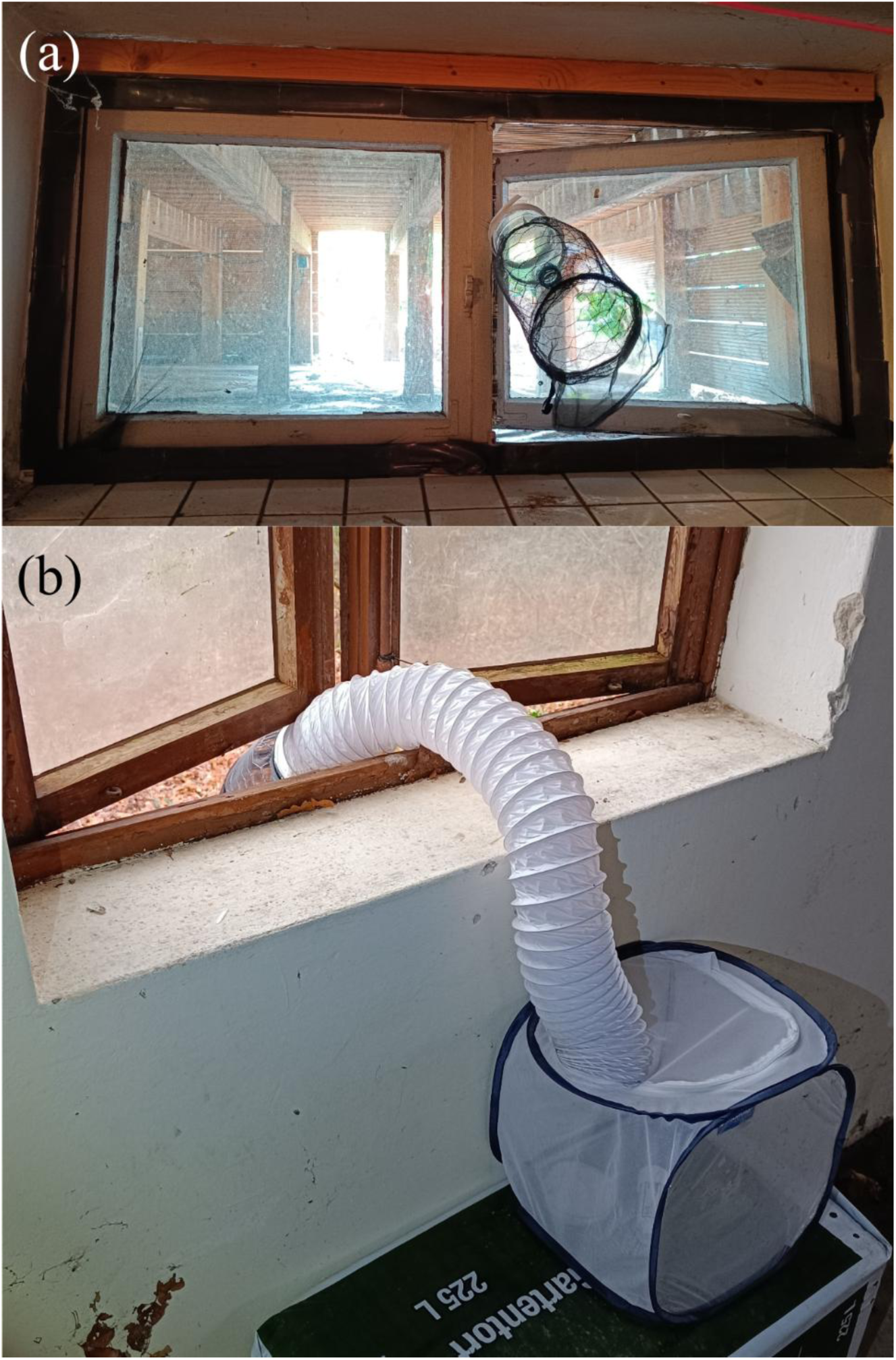
Representative images of (a) an entry interception trap and (b) an exit trap.

Traps were inspected weekly until mid-November. Mosquitoes were collected using a handheld aspirator built following the model by GM Vazquez-Prokopec, WA Galvin, R Kelly and U Kitron [36]. The number of mosquitoes trapped was noted. Collected mosquitoes were subsequently released into an insect cage (AERARIUM, 30 cm × 30 cm × 30 cm, Bioform, Nürnberg, Germany) placed inside the overwintering sites. Each cage contained a Petri dish (⌀ 90 mm) and moist cotton to provide water for hydration. Indoor temperature and relative humidity inside the cages were recorded using HOBO Pro v2 U23-001A loggers (Onset Computer Corporation, Bourne, MA, USA) at hourly intervals. To capture external conditions, environmental temperature was monitored outside the sites, within a 10 m radius of the site entry points, using HOBO Pendant UA-002-64 loggers (Onset Computer Corporation, Bourne, MA, USA), also at hourly intervals.

During autumn and winter, insect cages were checked every two weeks to replace the moist cotton used for hydration. Dead specimens were collected for identification. Data loggers were replaced in mid-December to ensure sufficient battery capacity for continued data recording.

### Phase 2: Exiting overwintering sites

During spring, our goal was to assess the environmental drivers of mosquito exit timing. Exit from the overwintering sites was monitored weekly at all sites where mosquitoes had previously been trapped, between mid-February and mid-April in 2024 and 2025. Winter mortality was high in some sites. In the first year, when fewer than 20 specimens remained at a given site, additional mosquitoes found inside the overwintering structures—which had entered through unsealed openings such as doors opened by residents—were collected by hand aspiration. Surviving mosquitoes from sites with higher numbers were also relocated on the day of set up of the exit traps to those with lower abundance to ensure a sufficient sample size for the experiment. When specimens were relocated to a site, they were maintained in separate cages in order to preserve information on the microclimatic conditions associated with their original overwintering sites. In the second year, mortality was high overall, and we were forced to collect mosquitoes from a new location. All mosquitoes introduced originated from a single unmonitored donor site located in a rural area in Lower Saxony (Neufirrel/Uplengen). The donor site was an unheated, traditional farm potato cellar, offering cool, stable subterranean conditions. Because all introduced mosquitoes originated from this single site, they experienced similar overwintering conditions prior to relocation. Consequently, differences in subsequent exit timing among experimental sites primarily reflect the local environmental conditions experienced during the spring monitoring period rather than differences in prior overwintering history.

In this phase, the exit traps consisted of a flexible white PVC hose reinforced with a rigid steel wire spiral (PVC Flex Hose Series K-660, K-TECH-PRO, Traitsching, Germany), inserted into the insect cage inside the overwintering site, and secured in place using the cage zipper. The hose extended to the outside of the overwintering site, where the previously described custom-made mosquito trap was attached at its distal end (Figure 1b). Microclimatic data were collected as described above with loggers inside and outside the overwintering sites. Traps were inspected weekly, and mosquitoes were collected using the same procedure described for the previous phase. Collected mosquitoes were kept in the catch bags and transported to the laboratory where they were frozen at -18 °C.

### Mosquito identification

Specimens were kept on a cold pack and morphologically identified to the lowest level of taxonomic certainty, using the morphological key by N Becker, D Petrić, M Zgomba, C Boase, MB Madon, C Dahl and A Kaiser [37]. Identification was carried out using a LEICA MC170 HD binocular microscope (Leica Microsystems, Heerbrugg, Switzerland). *Culex pipiens* s.s./*Cx. torrentium* specimens were typed using multiplex real-time PCR as described by M Rudolf, C Czajka, J Borstler, C Melaun, H Jost, H von Thien, M Badusche, N Becker, J Schmidt-Chanasit, A Kruger, et al. [38].

### Wing size measurements

Before molecular species identification, the wings of the mosquitoes were removed, mounted in Euparal (Carl Roth, Karlsruhe, Germany), and subsequently photographed using either an Olympus SZ61 stereomicroscope (Olympus, Tokyo, Japan) for the mosquitoes collected in 2024 or an Olympus SZX10 (Olympus, Tokyo, Japan) for the mosquitoes collected in 2025. Both stereomicroscopes were equipped with the same camera (Olympus DP23, Olympus, Tokyo, Japan). The centroid size of each wing was calculated based on 17 homologous landmarks coordinates which were collected by means of ITHILDIN, a web-based tool for automatic landmark detection in Diptera wings (https://ithildin.bnitm.de/, [39]). The resulting centroid sizes were averaged across the two wings of each specimen and used as a dimensionless relative proxy for wing size.

### Statistical analysis

Due to the experimental design, species identification was only possible on specimens that were recovered after successfully exiting the overwintering site. Consequently, species-level identification was not available for mosquitoes entering the overwintering sites, and entry dynamics were analysed at the family level (Culicidae), preventing species-specific assessment of phenology. In contrast, statistical analyses of exit dynamics were restricted to *Cx. p. pipiens* (excluding all other taxa).

We defined three phenological variables based on the timing of mosquito activity: the day of the year (DOY) (1–365) at which 10%, 50%, and 90% of the cumulative number of mosquitoes had entered or exited the overwintering sites, hereafter referred to as Q_10_, Q_50_, and Q_90_. These metrics were calculated at the site level for each winter, resulting in a single observation per site and year. As this aggregation removed within-site replication, mixed-effects models were no longer necessary.

For the entry phase, predictors included land use, ALAN, and environmental temperature. Linear models were used to test associations between phenological variables and environmental predictors. For the environmental temperature, we derived three summary metrics for the time period between each collection event: the mean, the mean of the lowest 10% of values (P_10_), and the mean of the highest 10% of values (P_90_). Because environmental temperature naturally declines late in the year as autumn approaches, raw temperature measurements are strongly correlated with DOY. Analysing raw temperature values would make it impossible to determine whether mosquito activity is genuinely triggered by temperature cues or if the association is merely an artifact of seasonal timing. To untangle these confounded factors, we decided to rather test whether short-term environmental temperature anomalies directly influenced mosquito behaviour. To do this, we detrended the temperature data. First, we fitted linear models for each environmental temperature metric (mean, P_10_, and P_90_) as function of DOY. This regression defined the expected seasonal baseline, i.e. the predictable decrease in environmental temperature as autumn progresses. We then extracted the residuals, which represent temperature deviations from this seasonal trend (that is, whether a day was unusually warm or cool relative to the seasonal expectation). Finally, these residuals were then used to assess whether environmental temperature deviations affected the timing of mosquito entry (Q_10_, Q_50_, and Q_90_). The most informative predictor among these metrics was subsequently selected for analysis using Akaike’s Information Criterion (AIC). To further investigate potential nonlinear relationships between environmental temperature and mosquito phenology, generalized additive models (GAMs) [40] were fitted. To account for land use specific responses, separate smooth terms were fitted for each land use category, while including it also as an additional parametric term. Model fit was evaluated using adjusted R² and deviance explained.

For the exit phase, predictors included land use, indoor temperature, and indoor relative humidity. For both microclimatic variables, three summary metrics were derived: the mean, P_10_, and P_90_. Linear models were used to assess the relationships between the phenological variables and (i) land use, (ii) microclimatic predictors, and (iii) their combined effects in additive models, and the most informative summary metric was selected based on AIC and used for subsequent analyses. To directly compare the effect of all analysed predictors on mosquito overwintering behaviour, model-predicted marginal effect sizes were calculated. For ALAN, environmental and indoor temperature metrics, and indoor relative humidity metrics, the total marginal effect was computed as the difference in predicted DOY (Δ DOY) across the full range of observed predictor values. For land use, Δ DOY represents the maximum absolute difference between categories.

In addition, fine-scale weekly entry and exit dynamics were analysed using cumulative degree-day frameworks. Preliminary models including DOY were also explored, but were excluded from the final analysis due to strong collinearity with cumulative degree-days (CDD), as both variables naturally co-varied with seasonal progression. For the entry phase, daily cooling degree-days (_cool_DD) were calculated across candidate base temperatures (*T*_base_ = 18, 16, 14 and 12 °C) using daily mean temperatures (*T*_mean_) from a local weather station of Germany’s national meteorological service (DWD, “Deutscher Wetterdienst”, weather station ID: 00691) via the formula max(0, *T*_base_ – *T*_mean_), summed over time to obtain cumulative cooling degree-days (C_cool_DD). The calculation of C_cool_DD used August 1^st^ as a fixed biofix date for both monitoring years. For the exit phase, daily CDD were calculated using four base temperatures (*T*_base_ = 0, 2, 5, and 8 °C) according to standard thermal accumulation protocols described by GS McMaster and WW Wilhelm [41], setting January 1^st^ as a fixed biofix date. Daily degree-days (DD; the thermal increment for a single day) were calculated using daily mean temperatures (*T*_mean_), computed as the average of 24-hourly logger readings, via the formula max(0, *T*_mean_ – *T*_base_), ensuring that values below the respective base temperatures were truncated to zero to prevent negative accumulation. CDD values were then calculated by cumulatively summing daily DD values over time. As sampling occurred weekly and indoor temperature data were recorded hourly, CDD values were linearly interpolated to match the precise sampling dates for each site. To test whether local environmental temperature was similarly predictive of exit timing, cumulative degree-days based on environmental temperatures were evaluated using the same framework. Note that the weekly exit analysis was restricted to the 2024 sampling period as in 2025 high overwintering mortality required the introduction of mosquitoes collected from an unmonitored site, preventing inference about site-specific thermal accumulation prior to emergence. For both entry and exit, the cumulative proportion of mosquitoes entering or exiting the overwintering sites at each sampling event (reflecting the ratio relative to the total number of individuals sampled in traps rather than absolute local population estimates) was modelled as a function of standardised C_cool_DD or CDD, respectively, using generalized linear mixed-effects models (GLMMs; Bates et al. 2015. The models were fitted with binomial error distribution and ‘logit’ link function, including site as a random term. Land use type was initially included as a categorical predictor, along with its interaction with C_cool_DD or CDD. Model simplification was performed by sequentially removing non-significant terms. Models across candidate base temperatures (with and without land use effects) were evaluated and compared using AIC to select the most parsimonious thermal predictor for each phase.

We tested whether wing size in *Cx. p. pipiens* influenced the overwintering survival and exit timing from their overwintering sites in two separate statistical models. Overwintering survival, expressed as binary variable, was analysed in a GLMM with a binomial error distribution as a function of the standardized wing centroid size [42, 43]. To account for variation among study sites and study year, both were combined and included as random factor in the GLMM. A linear mixed model (LMM) was used to analyse the relationship between wing size and exiting timing [42, 43]. This LMM excluded the specimens that died during winter and only included the surviving *Cx. p. pipiens*, which were collected in the exit traps. Exit time per specimen, measured as DOY, was used as response with the corresponding standardized wing centroid size as fixed effect. Study site and study year were combined and included as random factor.

All statistical analyses were performed using ‘R’ (Version 4.0.5) [44] and the interface ‘RStudio’ [45]. The significance of predictors was assessed using Type II analysis of variance implemented in the ‘car’ package [46]. Post hoc pairwise comparisons among groups were conducted using Tukey-adjusted estimated marginal means implemented in the ‘emmeans’ package [47]. GAMs were fitted using the ‘mgcv’ package [40]. GLMMs were fitted with the ‘lme4’ package [42]. To summarise and visualise the main drivers of mosquito overwintering behaviour, model-predicted marginal effect sizes (Δ DOY) and 95% confidence intervals across predictor ranges were extracted using the ‘ggeffects’ package [48]. Finally, the ‘ggplot2’ [49] and ‘patchwork’ [50] packages were used for visualisation.

## Results

### Phase 1: Entry into overwintering sites

A total of 269 and 246 mosquitoes were captured in entry traps in autumn of 2023 and 2024, respectively, and were used to model entry phenology. Overall, environmental drivers were associated with mosquito entry timing into overwintering sites, with distinct effects observed across early (Q_10_), median (Q_50_), and late (Q_90_) phenological stages (Figure 2a). The timing of early mosquito entry, as measured by Q_10_, differed significantly among land use categories (Figure 3: *F* = 5.08, df = 2, *P* = 0.012, adjusted-*R*² = 0.18). Estimated marginal means indicated that semi-urban sites had the latest onset of entry (289 ± 6.5 days), followed by urban (275 ± 3.9 days) and rural sites (265 ± 4.1 days). Tukey-adjusted pairwise comparisons revealed that semi-urban sites differed significantly from rural sites by 24.1 ± 7.7 days (*P* = 0.009), corresponding to an overall predicted marginal shift of 24.1 days (Δ DOY, Figure 2a), while urban sites did not differ significantly from either rural (*P* = 0.22) or semi-urban sites (*P* = 0.14). These results indicate that land use is associated with variation in the timing of mosquito entry into overwintering sites, with semi-urban environments showing the latest entry. However, for both Q_50_ and Q_90_, representing median and late mosquito entry, there were no significant differences among land use categories (*P* = 0.290 and *P* = 0.081, respectively).

**Figure 2.**
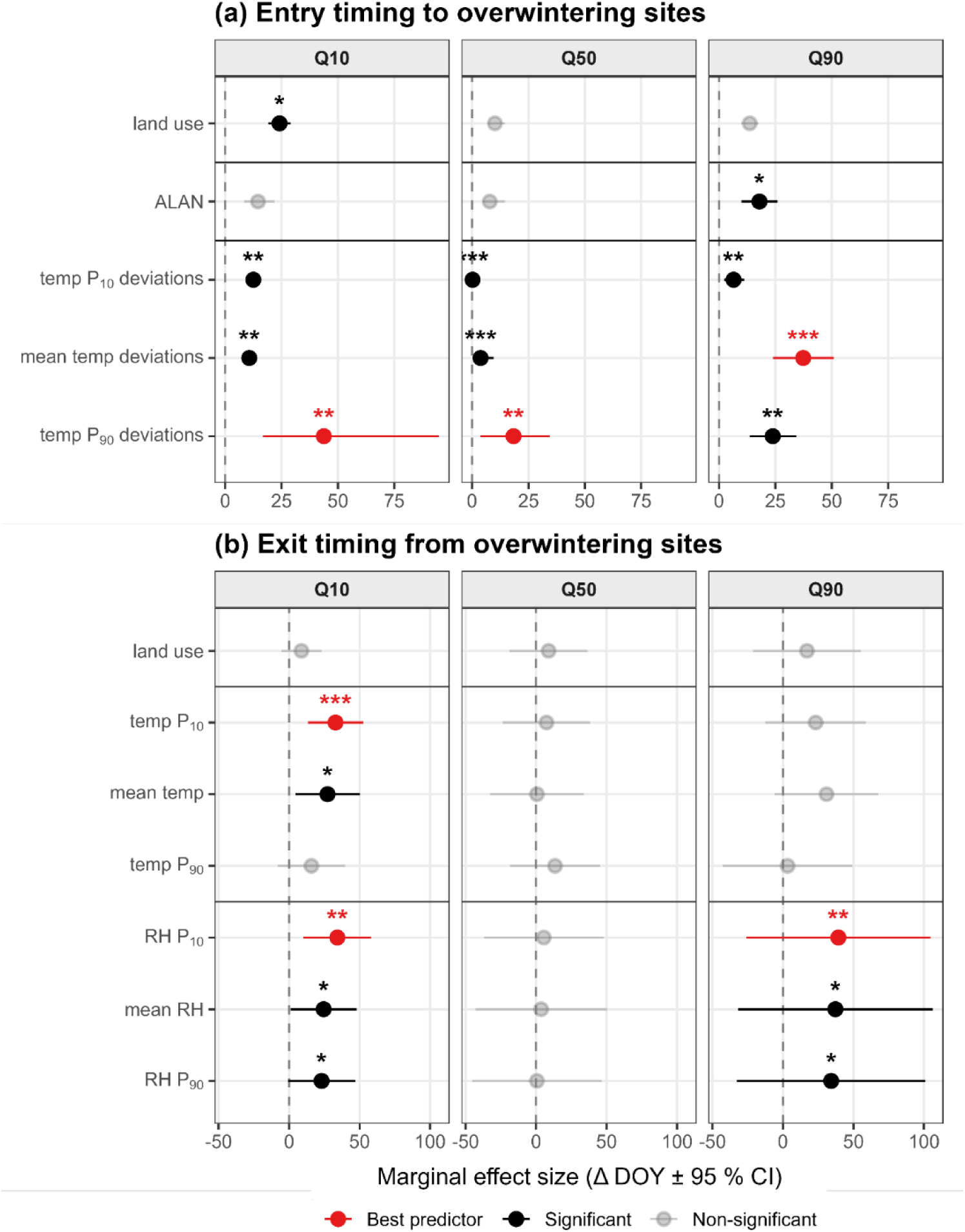
Summary of the marginal effect sizes (Δ DOY ± 95% CI) of environmental predictors associated with mosquito entry timing (a) and exit timing (b) across phenological phases (Q_10_ = early, Q_50_ = median, Q_90_ = late). Points represent total predicted effect sizes derived from model-averaged marginal predictions, with horizontal bars indicating 95% confidence intervals. Predictor categories are color-coded based on statistical performance: black = significant predictors (*P* < 0.05); grey = non-significant; red = best supported predictor within the variable category, based on AIC values when more than one metric was analysed per variable. Asterisks represent statistical significance levels (* *P* < 0.05, ** *P* < 0.01, *** *P* < 0.001). The vertical dashed line at zero indicates no marginal effect. Environmental predictors include land use, artificial light at night (ALAN), temperature deviations (temp deviations) metrics, temperature (temp) metrics, and relative humidity (RH) metrics. P_10_ = the mean of the lowest 10% values; P_90_ = the mean of the highest 10% values.

**Figure 3.**
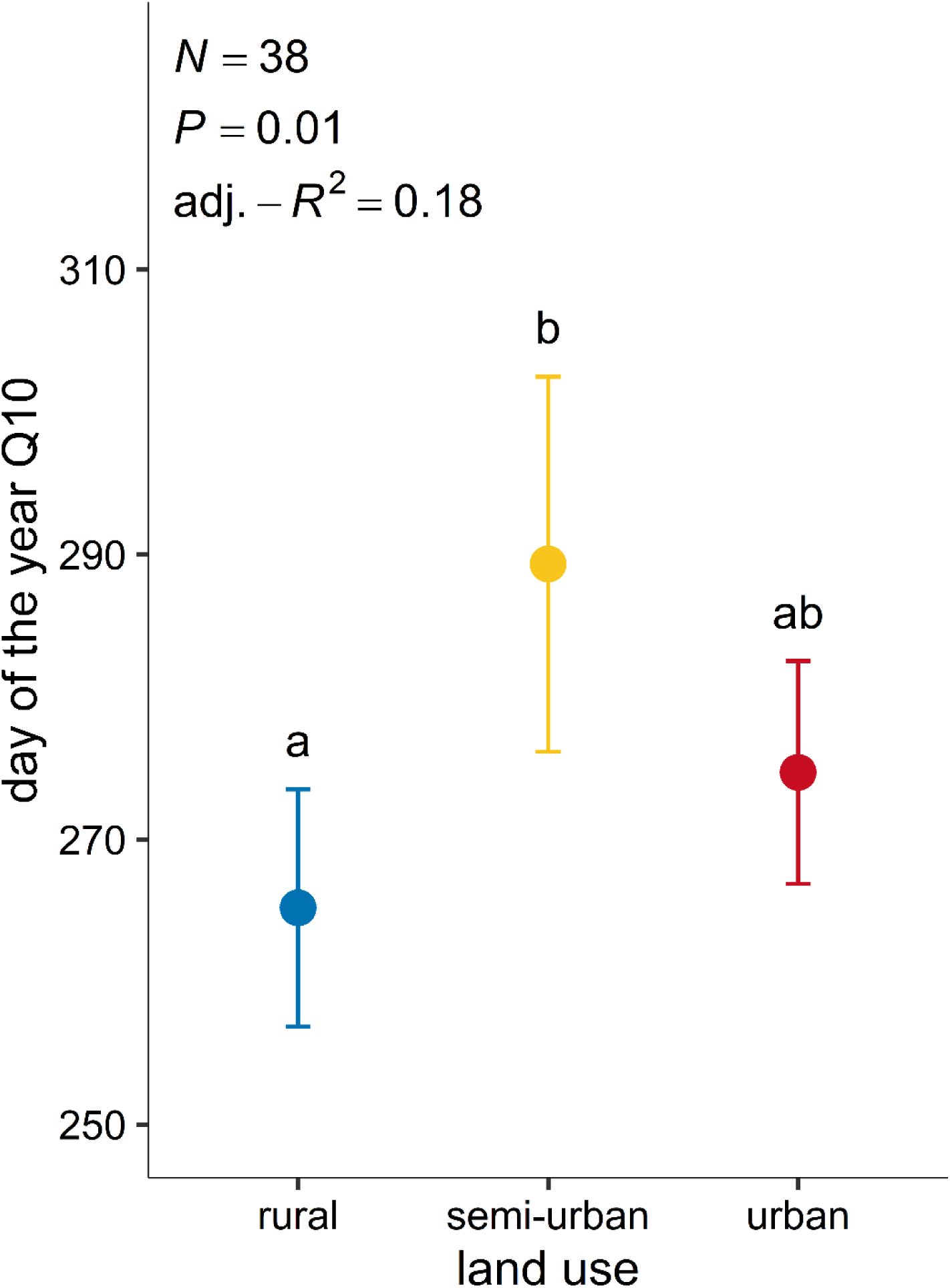
Estimated marginal means of Q_10_ entry (day of year in which 10% of the cumulative number of mosquitoes had entered the overwintering sites) across land use categories. Points represent predicted values from linear models, with error bars indicating 95% confidence intervals. Letters indicate Tukey-adjusted pairwise differences (a ≠ b).

For the early (Q_10_) and median (Q_50_) entry, ALAN had no significant effect (*P* = 0.150 and *P* = 0.361, respectively). In contrast, for late entry (Q_90_), ALAN was a significant predictor (Figure 4), with higher levels of ALAN associated with a later entry into the overwintering sites (*F* = 4.46, df = 1, *P* = 0.042, adjusted-*R*² = 0.09, y = 4.41x + 303.03). Model predictions revealed a total marginal delay of 17.9 days (Δ DOY, Figure 2a) across the observed ALAN gradient, (corresponding to an unstandardised rate of 0.64 days per unit increase in ALAN, or 4.41 days per SD increase), suggesting that ALAN delays the entry to the overwintering sites for the final portion of the population.

**Figure 4.**
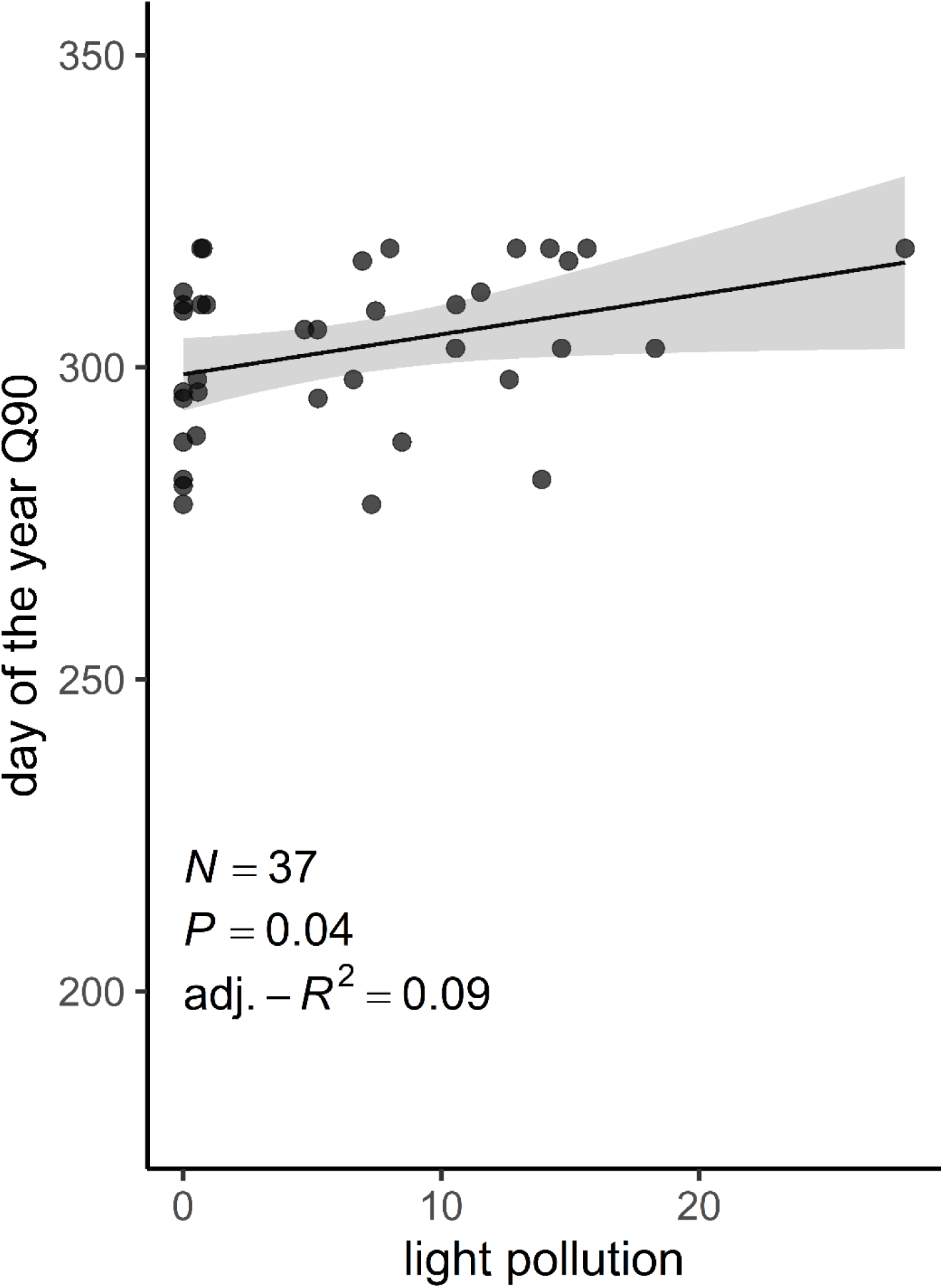
Predicted timing for Q_90_ entry (day of year in which 90% of the cumulative number of mosquitoes had entered the overwintering sites) as a function of artificial light at night (ALAN). Points show observed values, and the solid line represents the model prediction with the 95% confidence interval (shaded area).

Models revealed that environmental temperature deviations had a nonlinear effect on mosquito entry timing. For early entry (Q_10_), the better predictor was the mean of the highest 10% values (P_90_), i.e. environmental high-temperature deviations. The quadratic term was significant (*F* = 6.29, df = 1, *P* = 0.018), showing a total marginal effect size of 42.9 days (Δ DOY, Figure 2a) and showed a significant interaction with land use (*F* = 5.57, df = 1, *P* = 0.009), indicating that the environmental temperature–phenology relationship varied among land use categories. The GAMs supported these patterns and provided further insight into their shape (see Additional file 1: Figure S1): when using land use-specific smooth terms, the model explained 52.8% of the deviance (adjusted-*R*² = 0.43), with a significant nonlinear effect of environmental temperature only in semi-urban sites (edf = 2.05, *F* = 3.15, *P* = 0.025), and no significant relationship in rural (*P* = 0.507) or urban sites (*P* = 0.250). Median mosquito entry (Q_50_) showed similar patterns (see Additional file 2: Figure S2): the quadratic term for P_90_ deviations was significant (*F* = 8.06, *P* = 0.008) resulting in a total marginal effect size of 18.0 days (Δ DOY, Figure 2a) and its interaction with land use remained significant (*F* = 8.03, *P* = 0.002). GAMs again indicated that the nonlinear environmental temperature effect was restricted to semi-urban sites (edf = 2.34, *F* = 5.53, *P* = 0.003), with no significant effect in rural or urban habitats (adjusted-*R*² = 0.40). For late mosquito entry (Q_90_), mean environmental temperature was the better predictor. Our models revealed a strong nonlinear effect of environmental temperature deviations, (quadratic term: *F* = 18.00, *P* < 0.001), generating a total marginal effect size of 37.3 days (Δ DOY, Figure 2a), while the linear term was not significant (*P* = 0.432). Unlike Q_10_ and Q_50_, neither land use (*P* = 0.268) nor its interaction with environmental temperature (*P* = 0.907) was associated with entry timing, indicating a consistent environmental temperature–phenology relationship across land use categories. GAMs supported these findings, confirming a significant nonlinear environmental temperature effect (edf = 2.80, *F* = 5.17, *P* = 0.004) that explained 34% of the deviance (adjusted-*R*² = 0.34; Figure 5).

**Figure 5.**
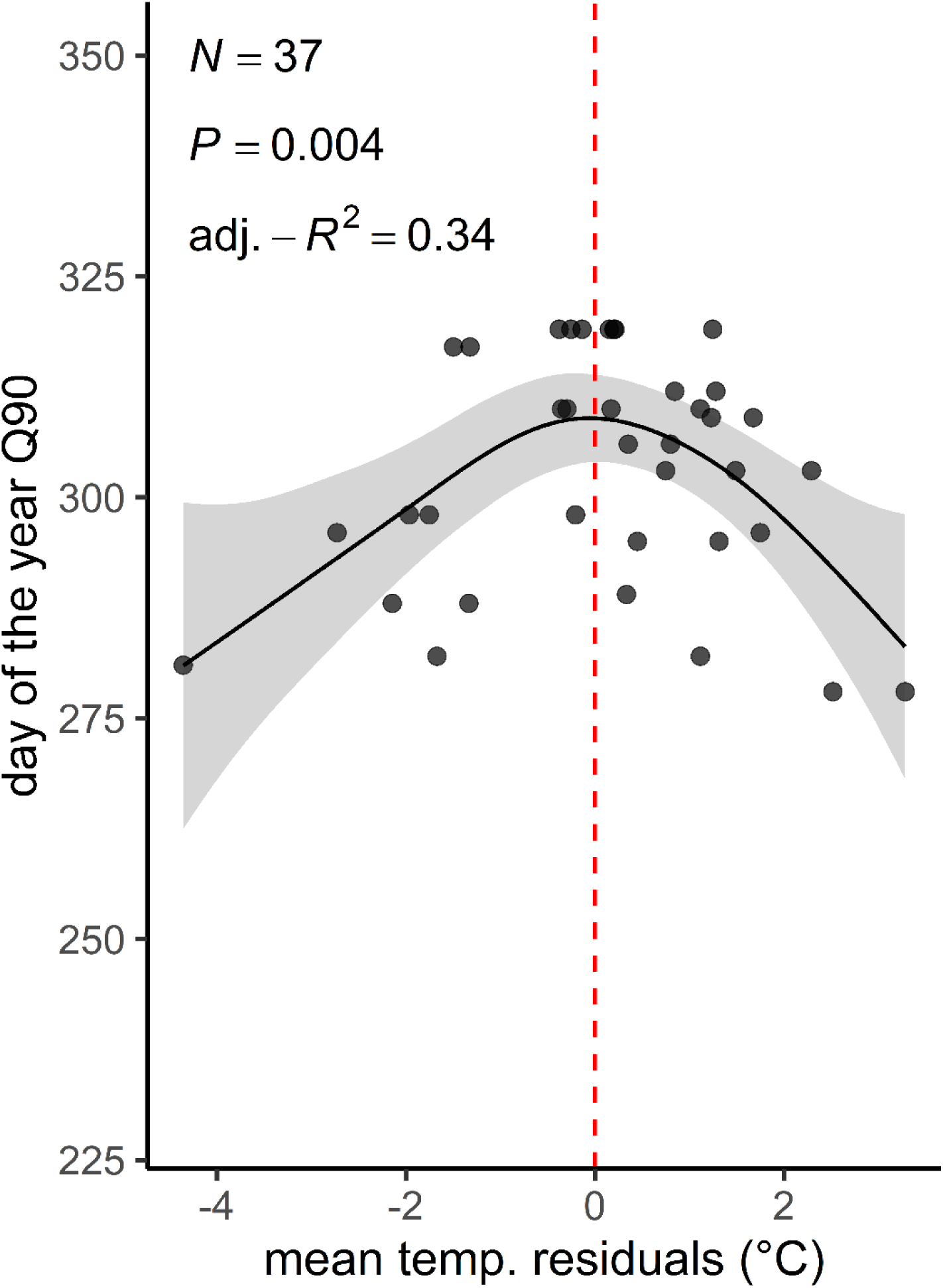
Nonlinear relationship between the detrended environmental mean temperature and the timing for Q_90_ entry (day of year in which 90% of the cumulative number of mosquitoes had entered the overwintering sites). Points show observed values (temperature residuals representing deviations from the seasonal trend). The solid line shows the fitted smooth from a generalized additive model (GAM), and the shaded area indicates the 95% confidence interval. The dashed vertical line marks zero residual temperature (i.e., expected seasonal conditions).

Finally, model comparison indicated that cumulative cooling degree-days (C_cool_DD) calculated with a base temperature of 18 °C provided the best fit (AIC = 818.1), outperforming models based on 16 °C (AIC = 870.4), 14 °C (AIC = 933.5), and 12 °C (AIC = 964.7). Model simplification showed that the interaction between land use and thermal accumulation was non-significant (*χ*^2^ = 2.16, df = 2, *P* = 0.339). However, the final additive model showed that both land use (*χ*^2^ = 6.84, df = 2, *P* = 0.033) and C_cool_DD (*χ*^2^ = 928.17, df = 1, *P* < 0.001) had a strong positive relationship with mosquito entry (see Additional file 3: Figure S3; marg.-*R*^2^ = 0.77, cond.*-R*^2^= 0.95), indicating that mosquitoes were increasingly likely to enter overwintering sites as cooling degree-days accumulated above the 18 °C threshold. Post hoc contrasts revealed that rural sites exhibited a significantly higher entry probability compared to semi-urban sites (OR = 4.21, *P* = 0.040), whereas semi-urban sites also differed significantly from urban sites (OR = 0.25, *P* = 0.045), with rural and urban sites showing no significant difference (*P* = 0.988).

### Phase 2: Exiting overwintering sites

Exit from the overwintering sites was monitored weekly between mid-February and mid-April in 2024 and 2025. Due to high overwintering mortality among the initial cohorts, experimental populations were supplemented prior to spring monitoring, resulting in a total study pool of 727 and 736 specimens across the two study years (Table 2). Across all processed specimens (*N* = 1,463), overall mortality across both study phases was 52.0% in winter 2023/2024 and 65.5% in winter 2024/2025. Out of all mosquitoes trapped, female *Culex* spp. represented 88.4% of the total capture, while *Cx. p. pipiens* represented 88.6% of the total *Cx.* spp. females. Other taxa collected included: *Aedes caspius*, *Anopheles maculipennis* complex, *Culiseta annulata*, *Culiseta* spp., and Culicidae that could not be further identified (Table 2). At the start of phase 2, 462 and 606 specimens were present in the overwintering sites in 2024 and 2025, with 349 and 254 specimens successfully exiting the overwintering sites, respectively (Table 2). For the subsequent analyses of the exit timing, we focused exclusively on female *Culex* spp. (comprising 91% of total emergence, or 549 specimens), among which *Cx. p. pipiens* represented 91% of the specimens (498 specimens).

**Table 2.** Entering and exiting mosquito counts per taxon across 2023/2024 and 2024/2025 study periods.

|  | 2023/2024 |  |  |  |  |  | 2024/2025 |  |  |  |  |  |
|---|---|---|---|---|---|---|---|---|---|---|---|---|
|  | entering |  |  | exiting |  |  | entering |  |  | exiting |  |  |
| Taxa | females | males | unknown | females | males | unknown | females | males | unknown | females | males | unknown |
| *Aedes caspius* | 0 | 0 | 0 | 0 | 0 | 0 | 1 | 0 | 0 | 0 | 0 | 0 |
| *Anopheles maculipennis* complex | 4 | 0 | 0 | 1 | 0 | 0 | 23 | 0 | 0 | 22 | 0 | 0 |
| *Culex pipiens* biotype *pipiens* | 527 | 6 | 28 | 289 | 0 | 0 | 619 | 1 | 1 | 209 | 1 | 0 |
| *Culex torrentium* | 3 | 0 | 0 | 2 | 0 | 0 | 0 | 0 | 0 | 0 | 0 | 0 |
| *Culex territans* | 0 | 0 | 0 | 0 | 0 | 0 | 1 | 0 | 0 | 0 | 0 | 0 |
| *Culex pipiens* biotype *pipiens* × *Cx. pipiens* biotype *molestus* | 0 | 0 | 0 | 0 | 0 | 0 | 2 | 0 | 0 | 1 | 0 | 0 |
| *Culex pipiens* s.s. | 0 | 0 | 0 | 0 | 0 | 0 | 18 | 0 | 0 | 0 | 0 | 0 |
| *Culex pipien*s s.s./*Cx. torrentium* | 64 | 0 | 0 | 31 | 0 | 0 | 60 | 0 | 0 | 17 | 0 | 0 |
| *Culex* sp. | 0 | 1 | 0 | 0 | 0 | 0 | 0 | 0 | 0 | 0 | 0 | 0 |
| *Culiseta annulata* | 34 | 4 | 0 | 12 | 0 | 0 | 9 | 0 | 0 | 4 | 0 | 0 |
| *Culiseta* sp. | 0 | 0 | 2 | 0 | 0 | 0 | 0 | 0 | 0 | 0 | 0 | 0 |
| Culicidae | 2 | 0 | 52 | 1 | 0 | 13 | 0 | 1 | 0 | 0 | 0 | 0 |
| Total | 634 | 11 | 82 | 336 | 0 | 13 | 733 | 2 | 1 | 253 | 1 | 0 |

Note that *Culex pipiens* s.s./*Cx. torrentium* represents mixed pools or specimens that could not be successfully differentiated via PCR amplification, while *Culex pipiens* biotype *pipiens* × *Cx. pipiens* biotype *molestus* denotes hybrid individuals between the two respective biotypes.

Overall, microclimatic conditions inside overwintering sites served as the primary drivers of mosquito exit timing, whereas land use showed no overall main effect across any phenological phase (Figure 2b) with no significant differences observed on early exit (Q_10_: *F* = 1.54, df = 2, *P* = 0.231), median exit (Q_50_: *F* = 0.51, df = 2, *P* = 0.607), or late exit (Q_90_: *F* = 1.10, df = 2, *P* = 0.392).

Indoor temperature was significantly associated with the timing of mosquito early exit (Q_10_), with the mean of the lowest 10% values (P_10_) providing the best model fit according to AIC. Higher minimum indoor temperatures were associated with earlier emergence (Figure 6: *F* = 14.00, df = 1, *P* < 0.001, adjusted-*R*^2^ = 0.28, y = -2.35x + 81.79), corresponding to an earlier exit timing by a marginal effect size of 27.7 days (Δ DOY, Figure 2b), or an unstandardised rate of 2.35 days earlier per 1°C increase in P_10_ indoor temperature. No significant relationship with any of the indoor temperature summary metrics was detected for median exit (Q_50_), nor late exit (Q_90_).

**Figure 6.**
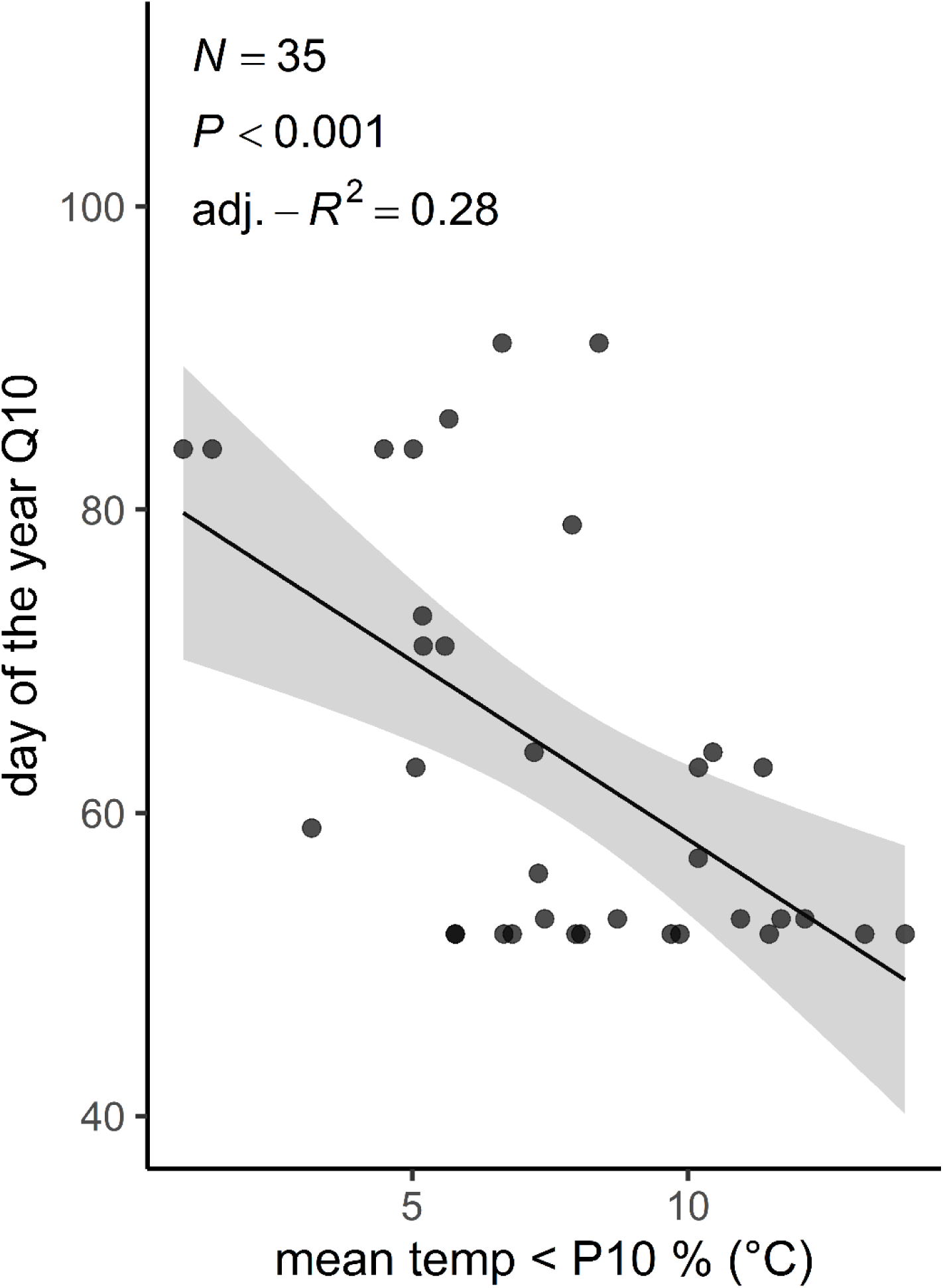
Relationship between the mean of the lowest 10% values (P_10_) of indoor temperature and the timing of Q_10_ exit (day of year in which 10% of the cumulative number of mosquitoes had exited the overwintering sites). Points show observed values, and the solid line represents the model prediction with the 95% confidence interval (shaded area).

Relative humidity was significantly associated with the timing of both early (Q_10_) and late (Q_90_) mosquito exit, with the mean of the lowest 10% values (P_10_) providing the best model fit in both cases according to AIC. However, relative humidity exhibited opposing relationships across these phases. For early exit (Q_10_), relative humidity (*F* = 12.15, df = 1, *P* = 0.001) and land use (*F* = 5.32, df = 2, *P* = 0.010) were significant predictors in an additive model (Figure 7, adjusted-*R*^2^ = 0.28), where higher humidity levels advanced early exit by 33.8 days (Δ DOY, Figure 2b), or 0.57 days per 1% increase in P_10_ relative humidity. Moreover, post hoc comparisons indicated that mosquitoes in urban sites exited significantly earlier than in rural sites (*P* = 0.011), while differences between rural and semi-urban sites were not significant (*P* = 0.999). The contrast between semi-urban and urban sites was not significant (*P* = 0.074). No significant association with relative humidity was detected for median exit (Q_50_) regardless of the summary metric considered. However, for late exit (Q_90_), lower relative humidity values (P_10_) was again a strong predictor (Figure 8; *F* = 14.47, df = 1, *P* = 0.007, adjusted-*R*^2^ = 0.63), with higher humidity levels corresponding to a delayed exit by 39.3 days (Δ DOY, Figure 2b).

**Figure 7.**
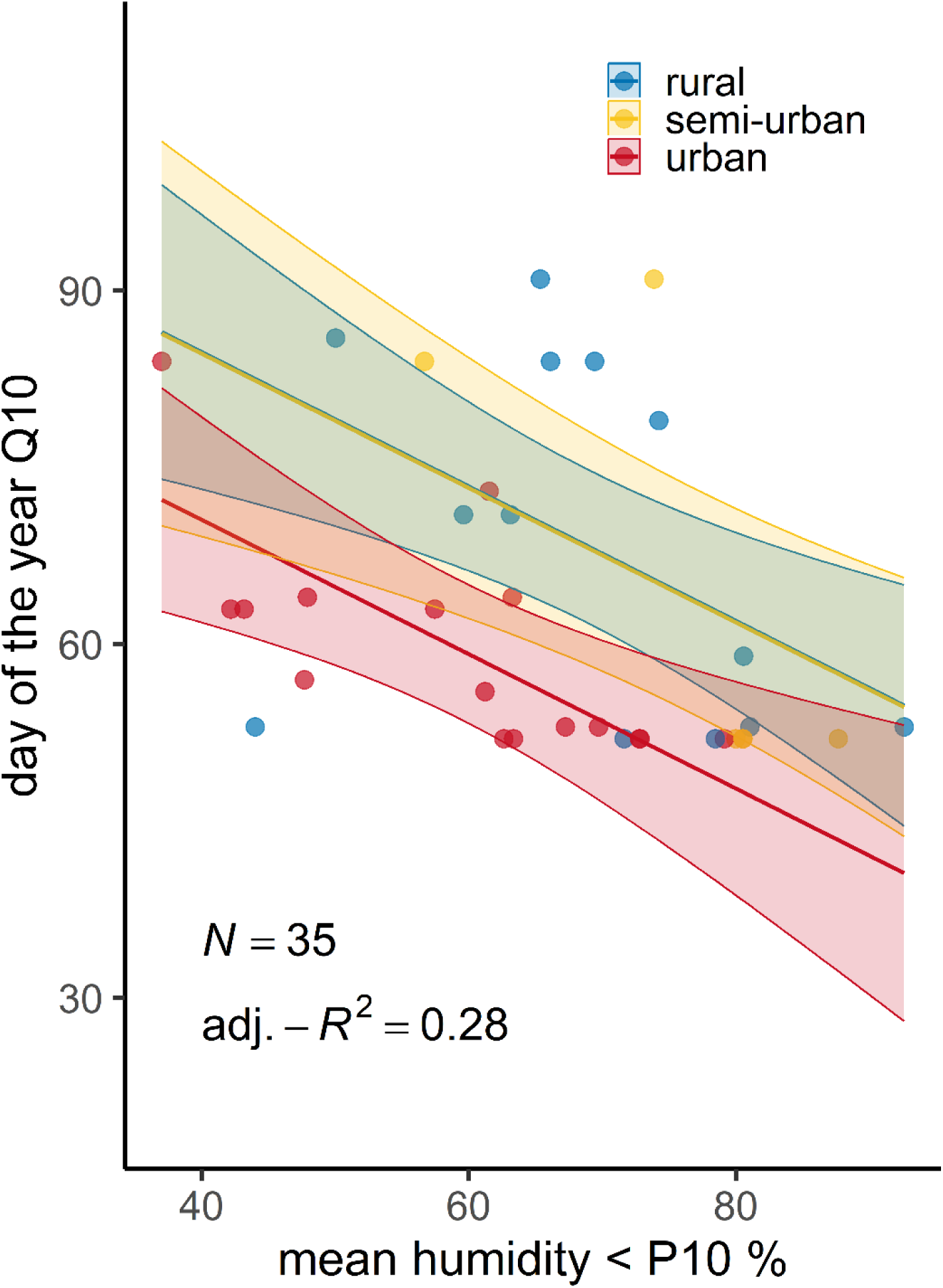
Relationship between the mean of the lowest 10% values (P_10_) of relative humidity (*F* = 12.99, df = 1, *P* = 0.001) inside the overwintering sites and the timing of Q_10_ exit (day of year in which 10% of the cumulative number of mosquitoes had exited the overwintering sites), across land use categories (*F* = 5.32, df = 2, *P* = 0.010). Points show observed values, and the solid line represents the model prediction with the 95% confidence interval (shaded area).

**Figure 8.**
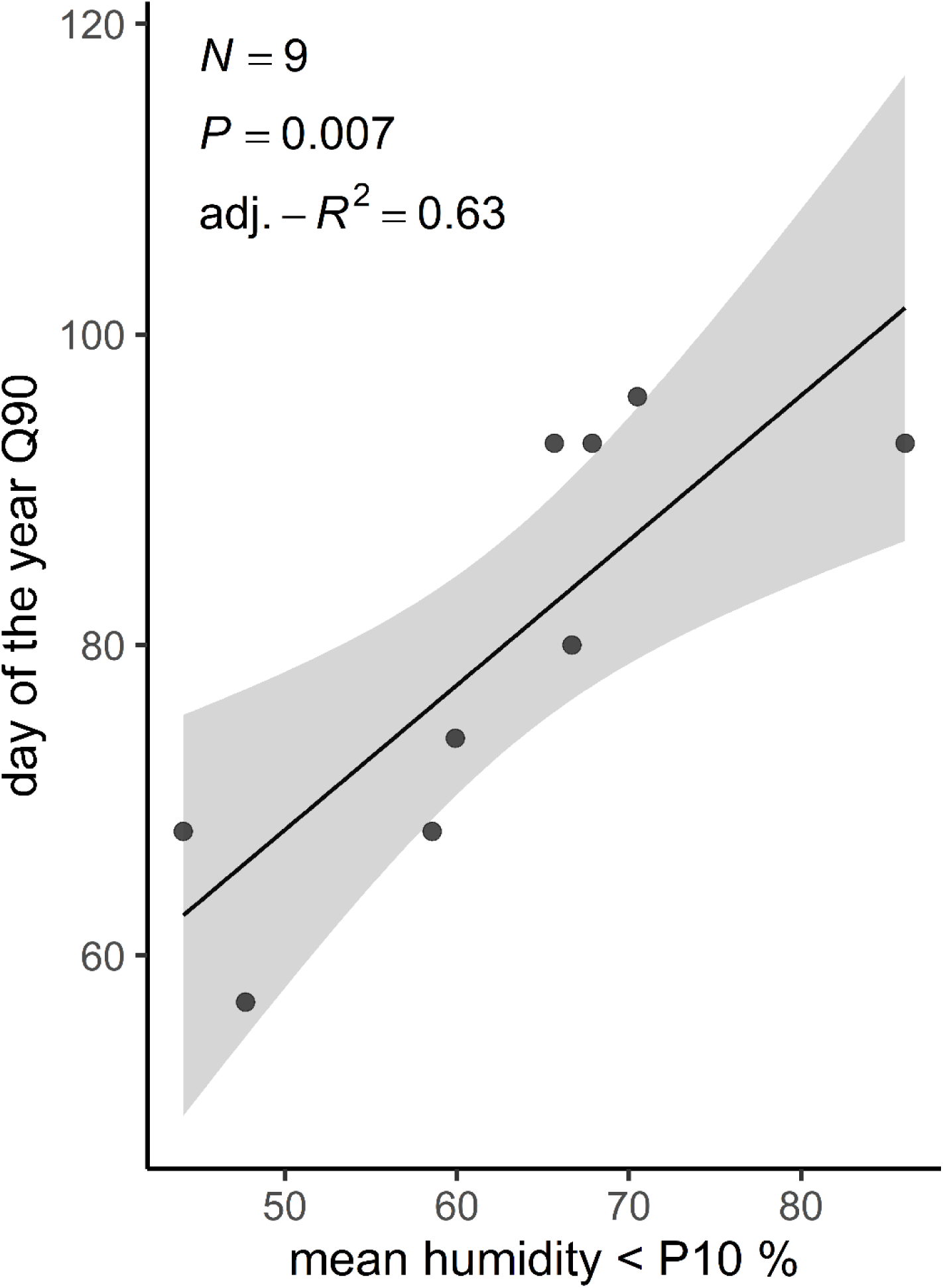
Relationship between the mean of the lowest 10% values (P_10_) of indoor relative humidity (*F* = 14.47, df = 1, *P* = 0.007) and the timing of Q_90_ exit (day of year in which 90% of the cumulative number of mosquitoes had exited the overwintering sites). Points show observed values, and the solid line represents the model prediction with the 95% confidence interval (shaded area).

Finally, model comparison indicated that CDD calculated with a base indoor temperature of 0 °C (DD_0_) provided the best fit (AIC = 911.82), outperforming models based on DD_2_ (AIC = 918.99), DD_5_ (AIC = 948.07) and DD_8_ (AIC = 1052.21). Model simplification showed that land use and its interaction with DD_0_ did not improve model fit, therefore both were excluded from the final model. The final model showed a strong positive relationship between CDD_0_ and mosquito exit (Figure 9; *χ*^2^ = 608.07, df = 1, *P* < 0.001, marg.-*R*^2^ = 0.63, cond.-*R*^2^ = 0.97), indicating that mosquitoes were more likely to leave as indoor temperature accumulated above the 0 °C threshold. Similarly, when evaluating cumulative degree-days based on local environmental temperatures (DD_0_, env. temp), models revealed a strong positive relationship with exit timing (see Additional file 4: Figure S4; *χ*^2^ = 683.59, df = 1, *P* < 0.001, marg.-*R*^2^ = 0.76, cond.-*R*^2^ = 0.95).

**Figure 9.**
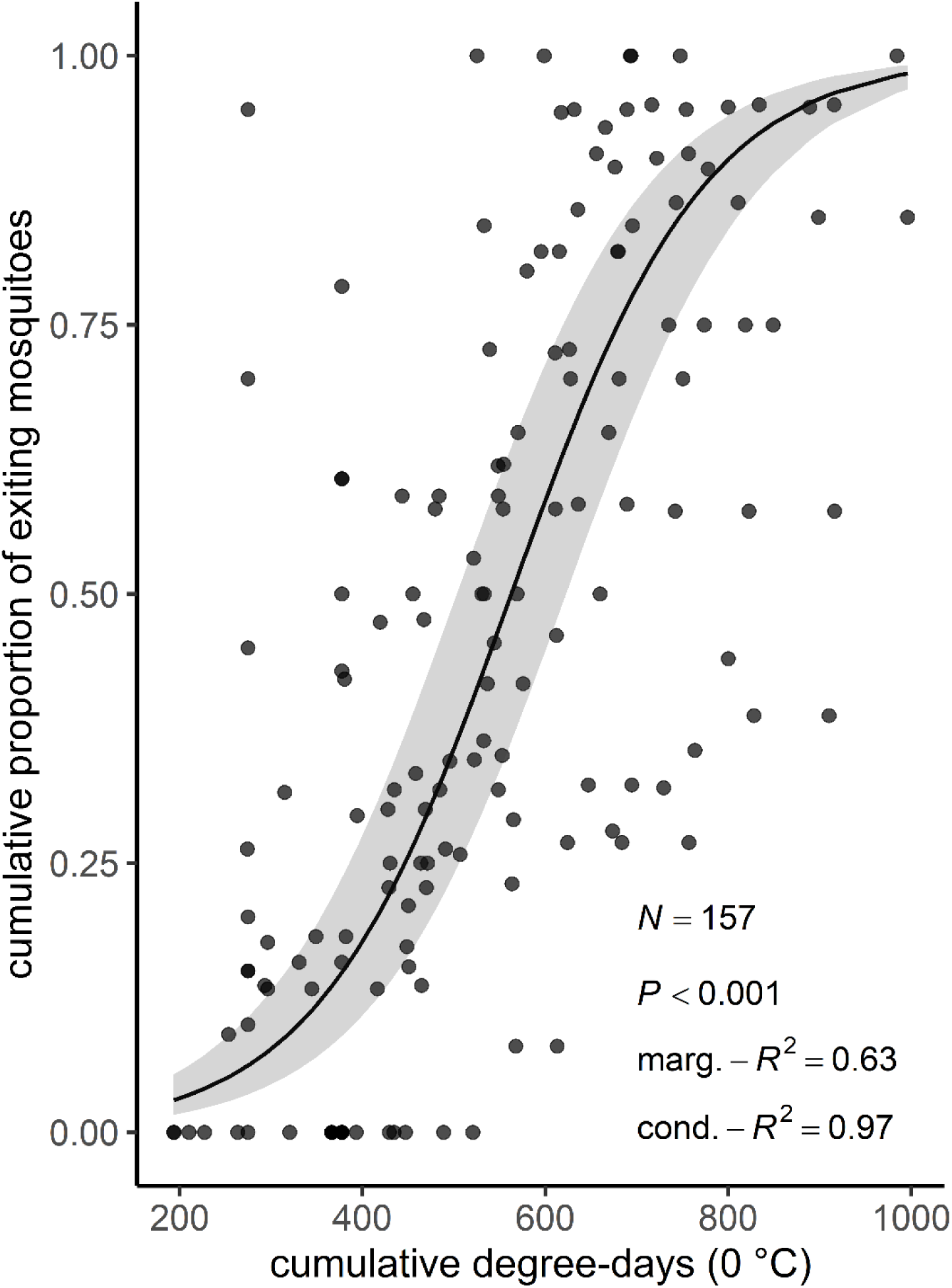
Relationship between cumulative degree-days calculated with a base indoor temperature of 0 °C (DD_0_) and the cumulative proportion of mosquitoes exiting the overwintering sites during the 2024 sampling period. Points show observed values, and the solid line represents the generalized linear mixed-effects model prediction with the 95% confidence interval (shaded area).

Due to damaged wings, the wing centroid sizes could only be measured in 805 of the 1,146 *Cx. p. pipiens* specimens. The wing size was not significantly associated with the overwintering survival in *Cx. p. pipiens* (GLMM, β = -0.07, SE = 0.11, z = -0.65, *P* = 0.52). However, the LMM, including only the surviving *Cx. p. pipiens*, showed a significant positive association between wing size and the exit DOY, indicating that *Cx. p. pipiens* individuals with larger wing size left the overwintering sites later in the year (β = 1.75, SE = 0.82, t = 2.14, df = 392.09, *P* = 0.033).

## Discussion

Our study shows that overwintering phenology in *Cx. p. pipiens* is shaped by a complex interplay of environmental and anthropogenic factors, with distinct drivers influencing entry into and exit from overwintering sites. Environmental temperature strongly modulated overwintering phenology, exhibiting non-linear effects during entry. In addition, land use and ALAN delayed overwintering entry, whereas warmer and more humid microclimates promoted earlier spring exit. Together, these findings highlight a high degree of behavioural plasticity, as adult mosquitoes dynamically adjust their entry and exit timing in response to localised anthropogenic and microclimatic cues.

### Phase 1: Entry into overwintering sites

Land use influenced the early phase of overwintering entry (Q_10_), with mosquitoes entering significantly later in semi-urban than in rural sites (as also illustrated by the overall cumulative trajectory in Additional file 3: Figure S3). Semi-urban habitats, represented by allotment gardens with heterogeneous vegetation structures, may provide abundant late-summer floral resources. An extended resource availability may allow females to continue sugar feeding and maximise lipid accumulation before seeking shelter, while habitats with scarce resources might lead to an earlier entry [9, 51]. However, land use effects disappeared later in the entry period. This shift suggests that seasonal cues such as declining environmental temperature and shortening photoperiods, ultimately override habitat-specific influences as the season progresses.

Although the observed effect size was moderate, ALAN was associated with delayed late entry into overwintering sites. A study by LR Fyie, MM Gardiner and ME Meuti [33] reported that exposing *Cx. p. pipiens* to ALAN altered diapause phenology by maintaining reproductive activity and fat accumulation simultaneously, resulting in lipid reserves that were lower than typical diapausing levels, yet higher than those of non-diapausing females. Our results align with these findings, suggesting that mosquitoes from urban environments associated with higher ALAN exposure remain active longer and enter overwintering shelters later in the season than those in rural environments.

Our findings reveal that mosquito overwintering entry is driven by non-linear thermal triggers, that shift from locally mediated, heterogeneous responses early in the season, to a uniform response by the end of the season. During early and median entry phases (Q_10_ and Q_50_), deviations from the environmental high-temperatures (P_90_) exhibited a distinct U-shaped relationship that advanced entry exclusively in semi-urban sites. One way to interpret this pattern is through the concept of fine scale thermal mosaics [52]. The allotment gardens representing our semi-urban sites are mostly open landscapes characterised by a high structural heterogeneity. These complex habitats create a microclimatic mosaic, where short-term environmental temperature changes are amplified, quickly becoming ecologically relevant [52, 53]. Within the semi-urban thermal mosaic, localised cold conditions may accelerate entry by signalling the rapid onset of winter, while warm anomalies might increase metabolic rate and energy consumption, potentially compromising overwintering survival [9, 54, 55]. However, given the limited sample size in semi-urban sites, the apparent non-linear responses should be interpreted with caution, as they may represent an artifact rather than an ecological driver.

In contrast to these heterogeneous semi-urban environments, neither urban nor rural sites responded to early-season anomalies. In urban areas, built structures likely dampen short-term environmental temperature fluctuations through the urban heat island effect, where thermal mass stores and releases heat predictably [56, 57]. Meanwhile, rural habitats lack intense anthropogenic modifications, and their local environmental temperature regimes may align more closely with regional seasonal climate patterns [56]. By late entry (Q_90_) however, local land use differences disappear; environmental temperature effects shift entirely to mean conditions, where both positive and negative anomalies advance entry. This once again suggests that later in the season, as environmental temperature declines and photoperiods shorten, broader seasonal environmental cues override the microclimatic variations provided by the local habitat structures.

When interpreting entry dynamics, it is important to distinguish between physiological diapause and post-emergence entry timing. While diapause in *Culex* mosquitoes is triggered during the aquatic larval stage via photoperiodic and thermal cues [9, 17, 58], our study focused specifically on adult behaviour. Post-emergence environmental conditions experienced by adults can modulate when diapausing females ultimately enter the overwintering sites.

Consequently, the observed variations in entry timing demonstrate post-emergence behavioural plasticity in response to immediate and localised environmental and anthropogenic drivers. However, three uncertainties remain regarding the precise mechanisms driving this timing. First, we did not study the larval stage, so we lack environmental data from the breeding sites where the adult mosquitoes developed potentially resulting in carry-over effects. Second, the time required for females to acquire sufficient lipid reserves through sugar feeding, and locate suitable overwintering sites is unknown. Third, macrohabitat variation in resource availability, dispersal distance, or shelter accessibility across land use categories may also influence entry timing. Therefore, observed shifts in entry timing likely reflect a combination of direct immature and adult responses to environmental drivers.

### Phase 2: Exiting overwintering sites

Land use alone did not significantly influence the overall timing of mosquito exit from overwintering sites across the full phenological range (Q_10_–Q_90_). This indicates that macrohabitat context plays a limited role in determining exit dynamics, which are instead driven primarily by microclimatic conditions in the overwintering site. Localised habitat effects emerged during the early exit. Early exit (Q_10_) was strongly advanced by higher minimum indoor temperatures (P_10_), reflecting the temperature-dependent nature of metabolic re-activation in *Culex* mosquitoes [9, 59]. Higher relative humidity similarly favoured early exit (Q_10_). Relative humidity has been shown to serve as a critical cue regulating dormancy in many insects [60, 61, 62, 63]. This is consistent with the physiological constraints of *Cx. p. pipiens* water balance: diapausing females rely on a heavily suppressed metabolism and cuticular hydrocarbons to limit water loss [64]. A higher relative humidity may therefore signal a low desiccation risk, prompting mosquitoes to safely re-activate their metabolism and terminate overwintering.

In addition to relative humidity, land use exerted a strong, additive effect, with mosquitoes in urban sites exhibiting earlier emergence than those in rural ones. Crucially, even when comparing sites at identical relative humidity levels, urban sites consistently promoted earlier emergence than rural sites. This effect points to urban drivers beyond moisture and aligns with the urban heat island effect, where elevated winter minimum temperatures accelerate thermal accumulation and metabolic re-activation within urban overwintering sites [56, 57]. In contrast, relative humidity had an opposing effect during the late exit phase (Q_90_), where higher relative humidity prolonged the emergence window regardless of land use category. In sites with higher relative humidity, reduced desiccation risk allows mosquitoes to persist longer before exiting. Conversely, in drier sites elevated desiccation risk likely forces late-persisting individuals to emerge earlier or face mortality, ultimately shortening the overall exit window.

Cumulative degree-days using a low developmental threshold (0 °C) provided the strongest predictor of exit timing (Figure 9), whereas higher thermal thresholds and land use did not improve model fit. This strong positive relationship indicates that overwintering termination is primarily driven by low-threshold thermal accumulation and temperature-dependent physiological processes. While a baseline threshold of 0 °C might seem low compared to the baseline used in other spring emergence, developmental or even transmission studies [59, 65, 66, 67, 68], it is well-supported in literature for modelling seasonal population dynamics of *Cx. restuans* and *Cx. pipiens* in northern regions [69]. Eco-physiological studies on Central European populations of *Cx. p. pipiens* suggest that prolonged exposure to 0 °C can induce severe latent cold injuries, dramatically reducing survival probabilities [22]. Because population mortality or any other post-diapause fitness measures were not quantified in this study, a 0 °C threshold best fit does not necessarily imply a lack of cold-induced fitness costs. Instead, it demonstrates that for the cohort of females that successfully withstand the winter, any microclimatic temperature above freezing allows metabolic activity in *Cx. p. pipiens*, confirming that mosquitoes in temperate-climate can process low-level thermal units throughout the coldest months to time their spring emergence. Furthermore, evaluating thermal accumulation based on local environmental temperatures revealed a similarly robust relationship with exit timing (Additional file 4: Figure S4).

We found evidence that larger *Cx. p. pipiens* specimens left the overwintering sites later in the season. Larger mosquitoes store more lipid reserves and may therefore be able to sustain overwintering for a longer period [18]. This later exit timing may provide a fitness advantage as they can avoid unfavourable early-season conditions, such as cold spells during spring. Nevertheless, we found no clear relationship between wing size and overwintering survival, which is in line with R Blom, J Spitzen, T de Haan and CJM Koenraadt [70], who continuously collected mosquitoes in several bunkers during two winter seasons and observed no change in wing size during the winters.

### Ecological and epidemiological implications

When considering the full annual lifecycle of *Cx. p. pipiens*, these findings reveal that urbanisation and microclimate variations results into a compressed overwintering window that shortens their total winter dormancy from both ends. In the autumn, anthropogenic modifications like ALAN and localised resources delay entry timing, extending mosquito activity late into the season. In the spring, the urban heat island effect interacts with a low developmental threshold advance exit timing. While this shortened dormancy poses severe bioenergetic risks by accelerating lipid depletion [23, 54], which might also have negative effects on the overwintering population, the surviving cohort may potentially extend the active season of vectors, amplifying the risk of enzootic arbovirus transmission prior to the summer peaks. Furthermore, because adult female *Culex* mosquitoes can successfully harbour arboviruses throughout the winter in Europe [14, 15], these altered overwintering dynamics facilitate early-season pathogen persistence. Ultimately, our results demonstrate how short-term thermal anomalies and anthropogenic factors alter vector phenology, signalling how ongoing global change may expand the annual active season of temperate mosquitoes.

### Limitations

Several limitations should be considered when interpreting these findings. First, because this was an observational study, unmeasured site-specific variables ─such as alternative overwintering site density, surrounding sugar source availability, distances to be travelled─ may have influenced mosquito phenology. Second, high overwintering mortality required supplementation and relocation of mosquitoes during the exit phase. While donor-site mosquitoes shared identical conditions during winter, their exact pre-overwintering history during early autumn remains unknown. Consequently, the emergence data reflect primarily the environmental drivers of exit timing, rather than natural field survival rates. Additionally, molecular species-level identification was only achievable during spring exit, preventing species-specific analyses during autumn entry. Finally, the statistical power of certain comparisons was constrained by the study’s two-winter duration and uneven replication across the different land use categories.

## Conclusion

Our study demonstrates that, as observed in other temperate insects, overwintering phenology in *Cx. p. pipiens* is not driven by a singular seasonal cue. Instead, adult mosquitoes display high behavioural plasticity, adjusting their entry and exit timing in response to local microclimates and human-altered environments. While autumn entry is delayed by ALAN and localised thermal mosaics, spring exit is driven by low-threshold thermal accumulation and potentially accelerated by the urban heat island effect. These shifts reveal that overwintering phenology is highly plastic, effectively extending the active window of a key disease vector in human-altered landscapes. As global change drives urbanisation and climate warming across temperate Europe, integrating these complex microclimatic drivers into regional surveillance frameworks will become vital for predicting early-season pathogen persistence and optimising targeted vector management.

## Supplementary information

**Additional file 1: Figure S1.** Relationship between detrended P_90_ temperature deviations and mosquito early entry across land uses.

**Additional file 2: Figure S2.** Relationship between detrended P_90_ temperature deviations and mosquito median entry across land uses.

**Additional file 3: Figure S3.** Cumulative mosquito entry as a function of cooling degree-days across land uses.

**Additional file 4: Figure S4.** Cumulative mosquito exit as a function of environmental cumulative degree-days.

## Abbreviations

AIC: Akaike’s Information Criterion

ALAN: artificial light at night

C_cool_DD: cumulative cooling degree-days

CDD: cumulative degree-days

DD: daily degree-days

DOY: day of the year

GAM: generalized additive model

GLMM: generalized linear mixed model

LMM: linear mixed model

Q_10_, Q_50_, Q_90_: phenological variables based on the timing of mosquito activity: the day of the year (1–365) at which 10%, 50%, and 90% of the cumulative number of mosquitoes had entered or exited the overwintering sites

P_10_: the mean of the lowest 10% of values

P_90_: the mean of the highest 10% of values

## Acknowledgements

The authors want to thank volunteers Tim Schnörwangen for his support in the field setup, Annika Feldmann for her support in the field set up and monitoring, Rocío Domingo Orusco for her support with the morphological identification, Unchana Lange, Anna Wulff and Philine Grohmann-Falke for their help during laboratory work, and Esther Timmermann and Kristopher Nolte for the technical support.

## Availability of data and materials

The datasets used and analysed during the current study are available from the corresponding author on reasonable request.

## Authors’ contributions

Conceptualisation: EK, RL, and FGS; experimental set-up and data collection: CVdC; data analysis and figures: CVdC; wing size data collection and analysis: FGS; first draft: CVdC; writing and editing: all authors.

## Ethics approval and consent to participate

Not applicable.

## Consent for publication

Not applicable.

