## Supplemental materials for "Environmental and anthropogenic drivers of *Culex pipiens* biotype *pipiens* overwintering phenology"

### Additional file 1:

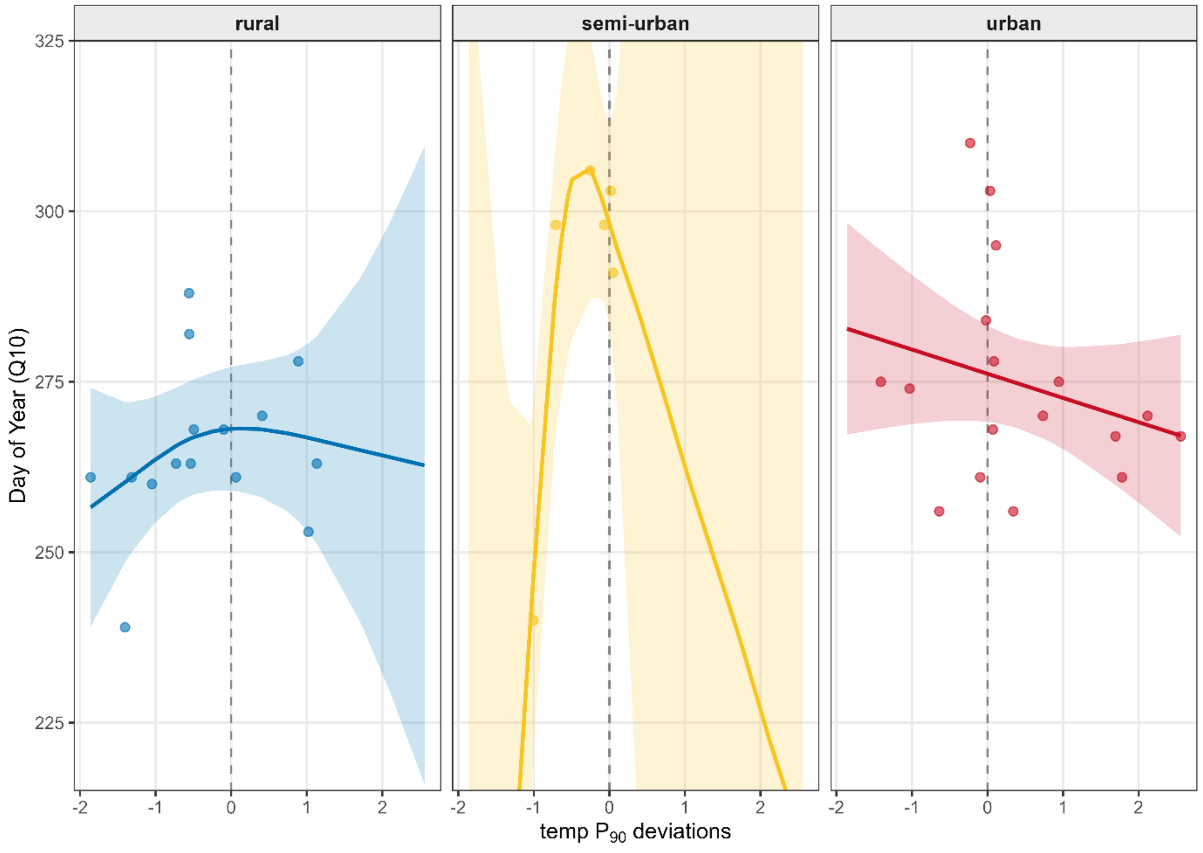


**Figure S1.** Relationship between detrended environmental $P_{90}$ temperature deviations (mean of the highest 10% values) and the timing of $Q_{10}$ entry (day of year in which 10% of the cumulative number of mosquitoes had entered the overwintering sites) across land use categories (rural, semi-urban, urban). Points show observed values, solid lines show land use-specific smooths from a generalized additive model (GAM), and shaded areas indicate 95% confidence intervals. The dashed vertical line marks zero residual temperature (i.e., expected seasonal conditions).

### Additional file 2:

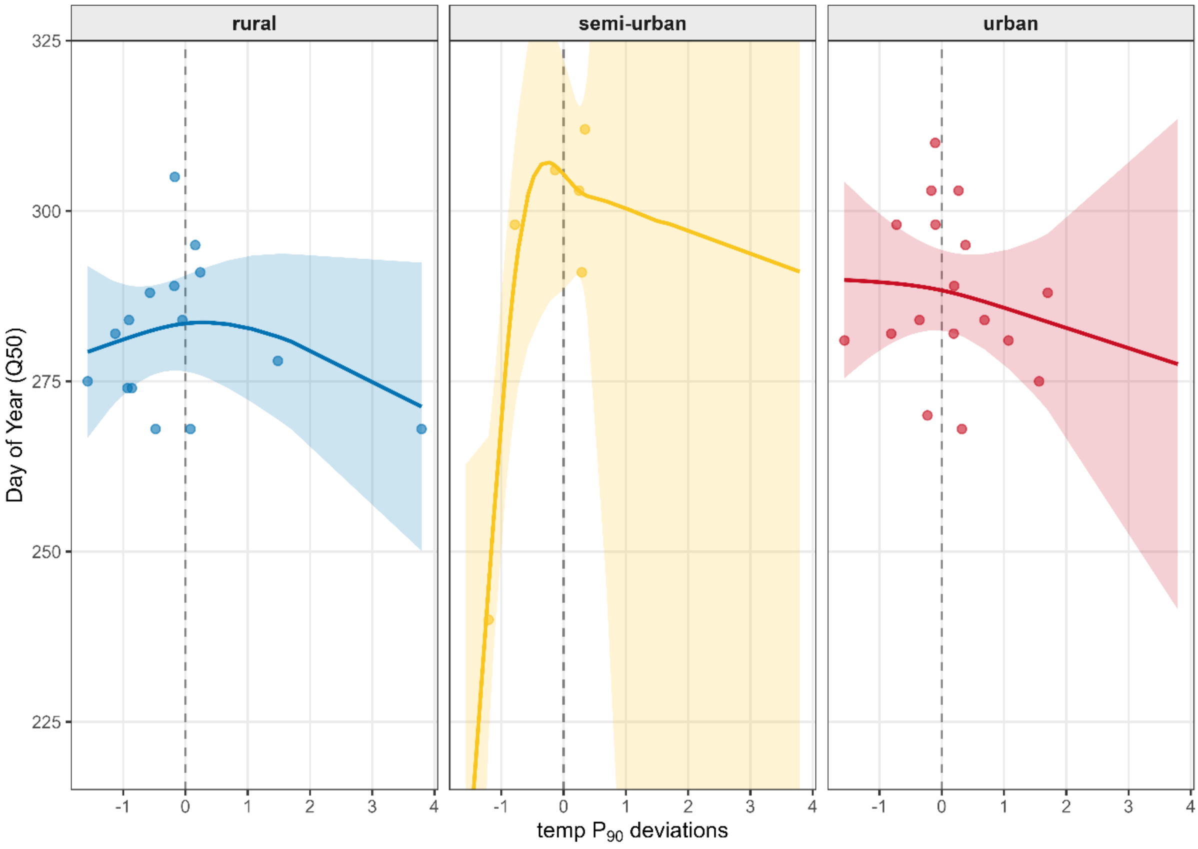


**Figure S2.** Relationship between detrended environmental $P_{90}$ temperature deviations (mean of the highest 10% values) and the timing of $Q_{50}$ entry (day of year in which 50% of the cumulative number of mosquitoes had entered the overwintering sites) across land use categories (rural, semi-urban, urban). Points show observed values, solid lines show land use-specific smooths from a generalized additive model (GAM), and shaded areas indicate 95% confidence intervals. The dashed vertical line marks zero residual temperature (i.e., expected seasonal conditions).

**Additional file 3:**

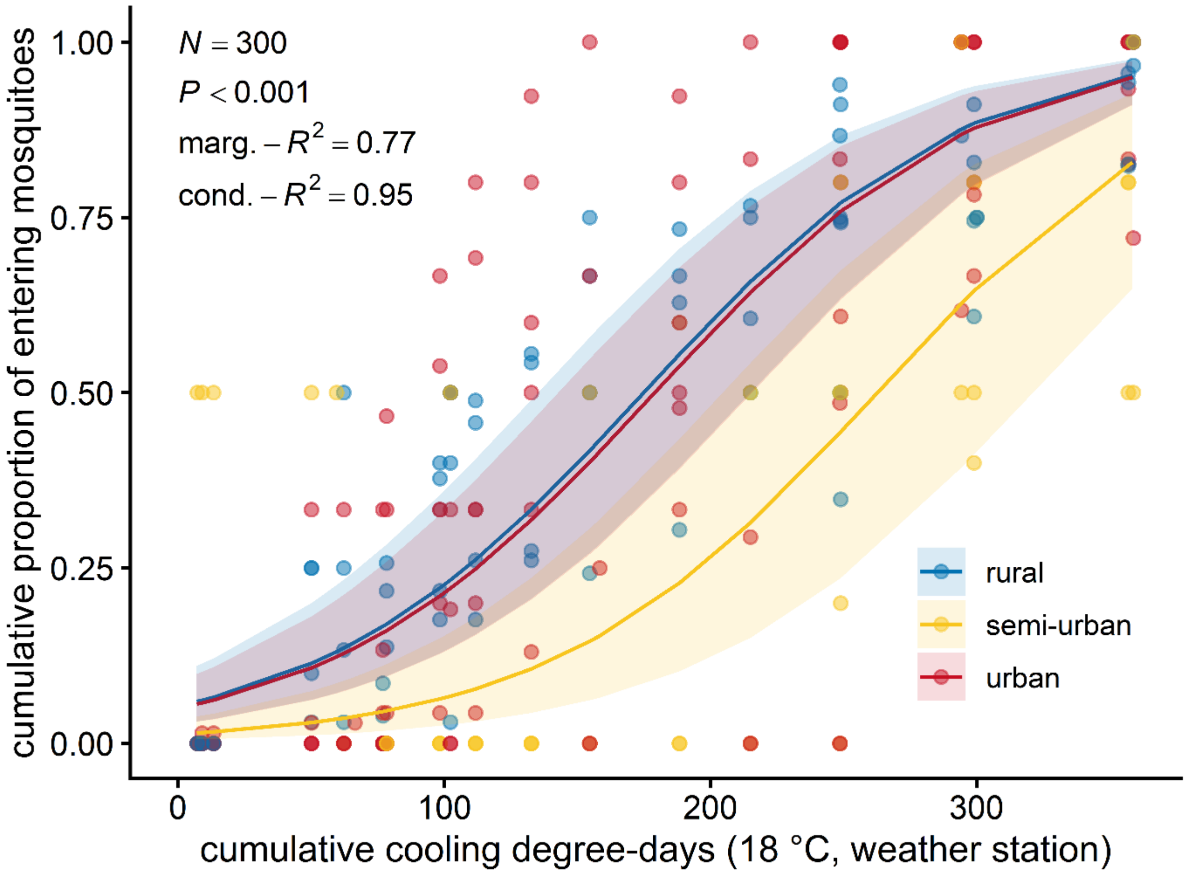


**Figure S3.** Cumulative proportion of mosquitoes entering the overwintering sites as a function of cumulative cooling degree-days ($C_{cool}DD$, calculated using a base 18 °C threshold from weather station data, starting from August 1st as biofix). Points show observed values across land use categories (rural in blue, semi-urban in yellow, and urban in red), and solid lines represent the generalized linear mixed-effects model predictions with the 95% confidence intervals (shaded areas).

**Additional file 4:**

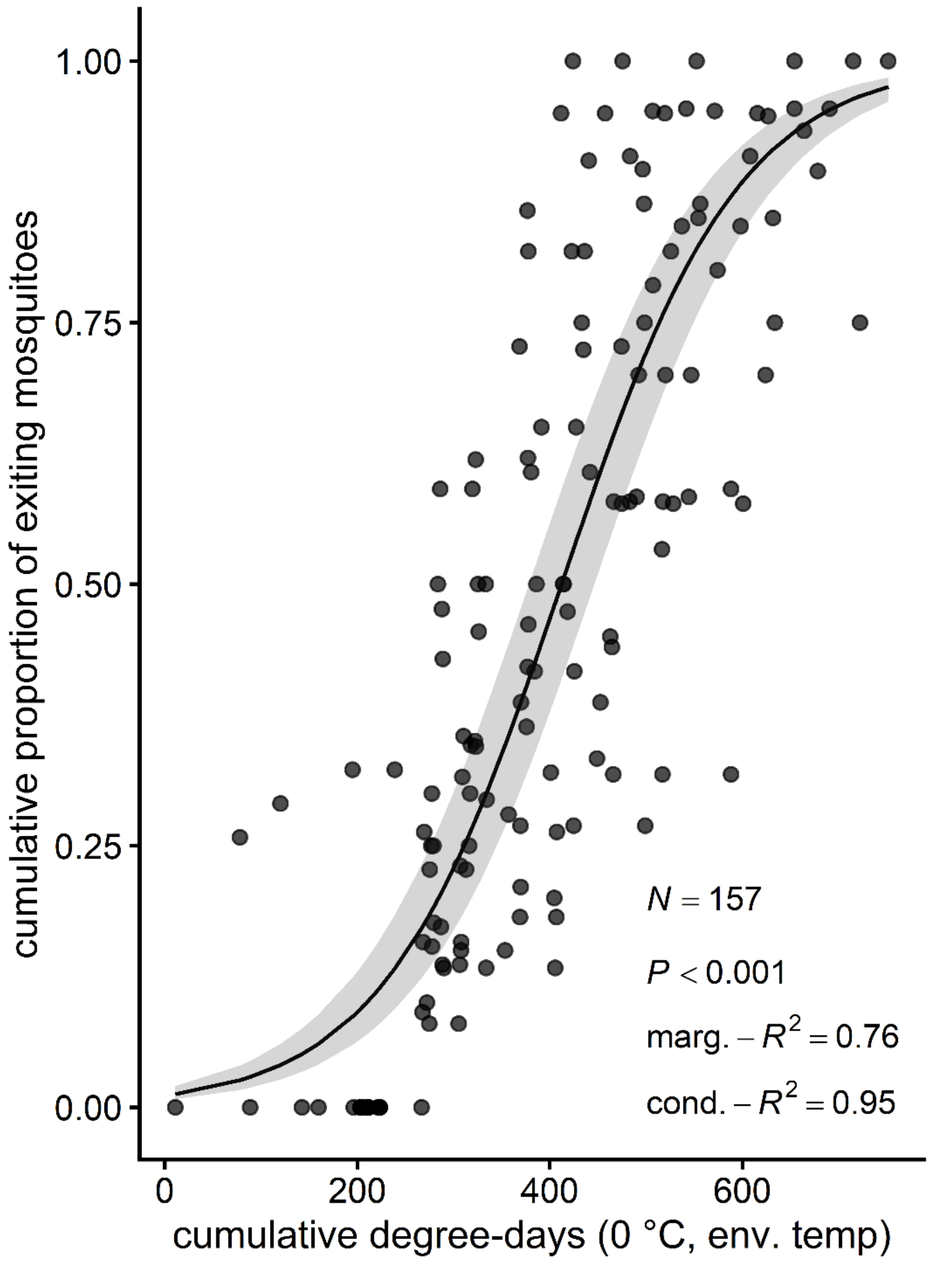


**Figure S4.** Cumulative proportion of exiting mosquitoes as a function of cumulative degree-days calculated using local environmental temperatures with a base of 0 °C ($DD_0$ env. temp) for the 2024 sampling period. Points show observed values, and the solid line represents the generalized linear mixed-effects model prediction with the 95% confidence interval (shaded area).
